# Engineered Peptides for Preventing Spy0128 Pilus Folding from *Streptococcus pyogenes*

**DOI:** 10.64898/2026.11.26.769537

**Authors:** Michelle Mendoza-Becerra, Tomás Hermosilla, Vicente Colarte, Sergio Canales, Benjamín Ulloa, J. Andrés Rivas Pardo

## Abstract

Intramolecular isopeptide bonds protect bacterial pilins from mechanical unfolding. Preventing isopeptide bond formation could compromise tissue colonization and infection. We previously demonstrated that a β-strand-derived peptide can interfere with bond formation in nascent Spy0128, the shaft pilin of *Streptococcus pyogenes*. Moreover, inserting an extra peptide into the protein’s core could disrupt proper folding, yielding extensible and mechanically labile proteins. Here, we explore computer-assisted isopeptide blocker peptides, designed specifically to interfere with the formation of isopeptide bonds present in the Spy0128 pilin protein. To accelerate our workflow, we implemented a computational pipeline that combines peptide library generation, structural preparation, and randomized molecular docking, all packed into a single software. Limiting peptide design to 10-residue ligands, we generated an initial library of 64 million peptides, which we reduced to fewer than 10 candidates after several iterations. Compared with the previously reported peptide, the binding energy changed from −7.6 kcal/mol to −9.0 kcal/mol. *In vitro* co-expression assays suggested peptide decoration with the designed peptide, similar to previous reports, but also confirmed abundant protein degradation, likely due to misfolding. Single-molecule magnetic tweezers assays confirm that the designed peptide interferes with the folding of Spy0128, yielding a fully extensible pilus protein.

## Introduction

Bacterial attachment requires adhesive structures that remain intact under the mechanical stresses imposed by the host. In *Streptococcus pyogenes*, the pilus consists of three main proteins: Spy0125, which anchors the entire filament to the cell wall; Spy130, also known as adhesin, mediating the binding of the bacterium to the epithelium; and Spy0128, which performs functions as the shaft domain, found in hundreds of repeated copies along the filament. An intramolecular isopeptide bond in this Spy0128 protein prevents mechanical extension of the folded domains. Accumulated evidence shows that preventing or blocking isopeptide bond formation affects tissue colonization; thus, it has been identified as a target to weaken the shaft before assembly into a mature pilus.

Structurally, the Spy0128 protein contains two tandem immunoglobulin-like domains; however, only the C-terminal domain is related to the force pathway, due to the polymerization between Spy0128 subunits, which connect the C-terminus of Spy0128 with the Lys161 from the following subunit, yielding only a short section of less than 10 residues that preceed the C-terminal domain (see ***Figure 1***). In the C-terminal domain, an isopeptide bond connects Lys179 to Asn303 and diverts the force pathway away from most of the folded structure. This crosslink’s location explains why changing bond formation can strongly affect the domain’s mechanical response. Nevertheless, interfering with bond formation requires acting before the pilin folds. In the past, a strategy based on competition addressed this idea; addressing the problem away from searching reagent that cleaves an existing covalent bond, we implement a peptide that competes for formation of the isoppetide bond. However, the success of the intervention rate was limited reaching level of peptide decoration close to 20%.

**Figure 1.**
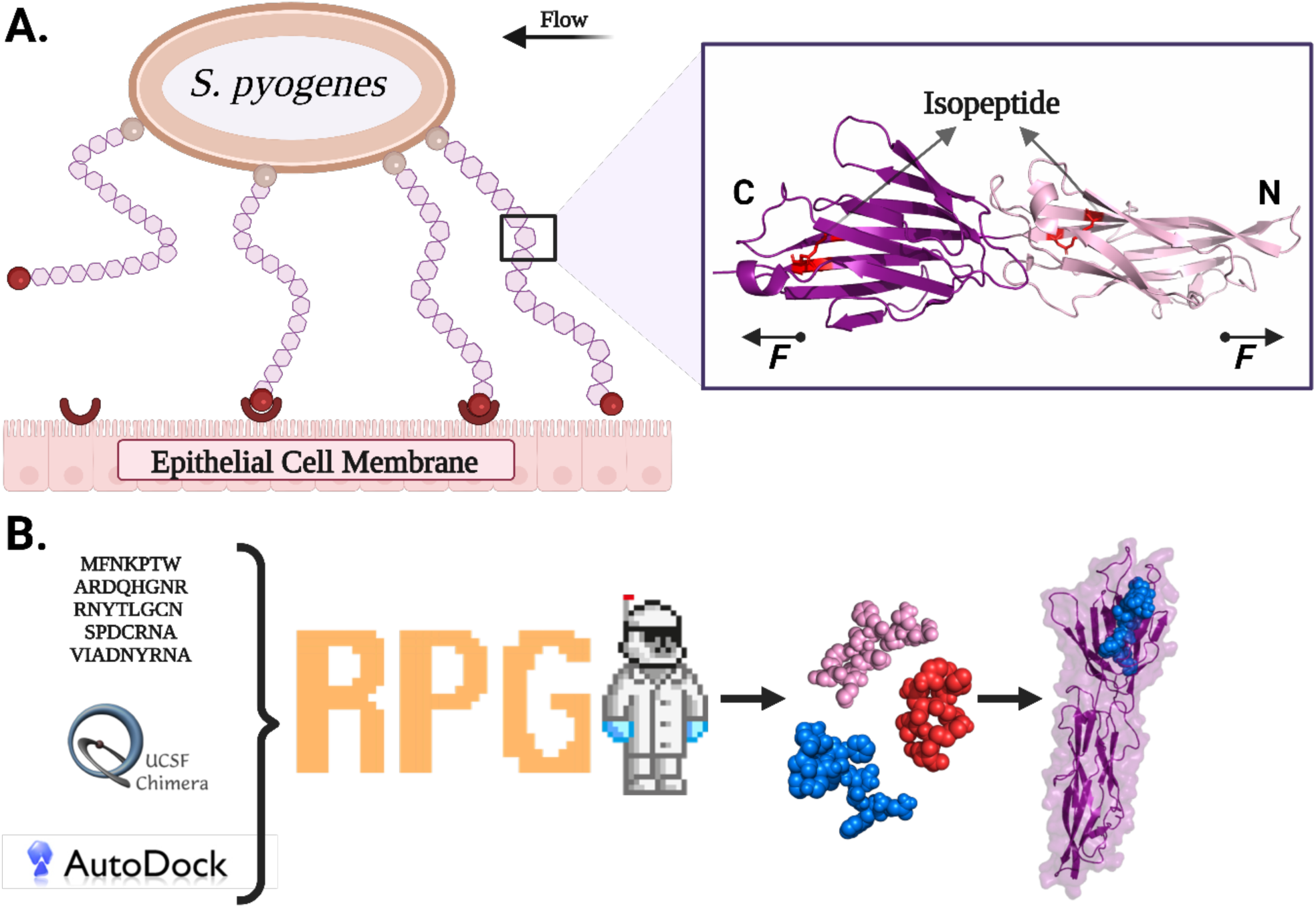
Computational strategy for selecting peptides directed at Spy0128 isopeptide bond formation A. S. pyogenes adheres to host surfaces through pili containing repeated Spy0128 shaft subunits. The intramolecular isopeptide bonds of Spy0128 prevent mechanical extension of its folded domains; the proposed intervention targets formation of the C-terminal Lys179–Asn303 bond during pilin folding [1]. B. RPG connects peptide generation, three-dimensional structure preparation, and docking against the target. The workflow is used to prioritize sequences for biochemical evaluation. The schematic illustrates the design rationale; it does not represent an adhesion or mechanical measurement performed in this study.

Sequential expression of the peptide derived from β-strand 11 of Spy0128 can be used as a perfect competitor of the natural β-strand; thus, interfering during early stage of the protein folding, is posible to prevent the intramolecular bond. The installation of the peptide in the middle of the molecule not only prevents the formation of the bond, but also has consequences for the mechanical stability of Spy0128, which turns into a fully extensible and mechanically labile protein.

Aiming to develop a platform for easy searching of isopeptide-blocked peptides, we implemented a workflow based on “random peptide generator” software, which allowed us to test virtually hundreds of peptides, select candidates, and later test them either through *in vitro* protein expression or single-molecule mechanics assays. Our results showed that computational screening contributed to selecting candidates for isopeptide blocker peptides; nonetheless, the effect of the peptide during the co-expression is hard to predict. C96 peptide triggers protein degradation and may be related to misfolding and toxicity for the bacterium.

## Material and Methods

### Peptide generation and Molecular docking

We implemented a workflow for computational prediction of protein-peptide complexes. For this, we developed an application that lets users build a peptide library, convert peptides into 3D structures, and then perform molecular docking between selected peptides and the protein target. The workflow app, which we called *Random Peptide Generator* (RPG), combines PeptideBuilder to generate PDB files, Open Babel to prepare the PDB files, and AutoDock Vina/Vinardo to conduct the molecular docking. Specifically, the user specified peptide length—limited to a total of 12 residues— and the amino acids allowed at each position. The generated sequences were written to a text file, establishing the peptide library, from which a selected number of peptides were converted into three-dimensional structures. Peptides were then docked against a predefined region of Spy0128, with a configuration file specifying the search region and peptide flexibility. Sequence identifiers and docking scores were displayed in the same app. A copy of the app is available on our GitHub.

For isopeptide-blocker peptide design, we fixed asparagine at position 4, which is needed to form the isopeptide bond with Lys179 included in the C-terminus of Spy0128. We substituted the other positions with the 20 standard amino acids. As a reference, we used RPG with the control peptide *KDMTITFT**N**K*(KDAE) and calculated its binding energy. We set this peptide as the reference threshold; binding energy below this value indicates a better isopeptide-blocker peptide, so we selected it for in vitro assays.

### Spy0128 Protein and Peptide Constructs

Because one aim is to test the peptides’ ability to prevent isopeptide bond formation on Spy0128, yielding an extensible protein molecule, we used mechanical reporters flanking Spy0128. Normally, intact Spy0128 is inextensible; therefore, no extension should be collected in the single-molecule mechanical assays, which is why I27 titin domains are included within the protein construct. I27 domain is a widely characterized protein, with extension and force well known; therefore, if I27 events are present in the recording, the force was also applied to the Spy0128. Thus, we cloned two copies of Spy0128 flanked by the I27 titin domain, increasing the chances of capturing Spy0128 in a single molecule. The polyprotein I27-Spy0128-I27-Spy0128-I27 was cloned into pFN18a vector, including a HaloTag and AviTag in the N-terminus and C-terminus, respectively. These two Tags allow protein immobilization, required for single-molecule force spectroscopy.

We designed and assayed a total of three peptides: *KDMTITFT**N**K(KDAE),* named the N14 peptide, previously proven as an isopeptide blocker; *QWG**N**MFNKWK,* referred to as the C96 peptide, specifically designed to interfere with Spy0128; and *EIK**N**IKQ*, a control peptide named D3, originally designed for targeting other pilus proteins, which was used as a specificity peptide control. We introduced the corresponding DNA sequence for each peptide into a custom-modified pBAD18 vector containing a copy of SNAP-Tag, a protein tag for protein labeling and isolation.

### Protein Expression and Purification

For protein expression, we initially tested several conditions to determine the optimal temperature and time to achieve the highest protein and peptide yield. Thus, we first tested optimal conditions for expressing either the Spy0128 polyprotein or the SNAP-tagged peptide. After confirming that both protein constructs were expressed without problems, we assayed coexpression conditions. Because the peptide will be competing to form the isopeptide bond with the emerging section of the Spy0128 polyprotein, we triggered the induction of the SNAP-tagged peptide, followed by the overexpression of Spy0128. By doing this, a high concentration is achieved within the bacteria before production of the Spy0128 polyprotein. Therefore, the expression was conducted in two stages: *i*) first, the bacteria were culture in minimal medium M9CA supplemented with 40 μg· mL−1 thymine, 2 mM magnesium sulfate, 0.1 mM calcium chloride, and 1.5% glycerol as carbon source until the culture reached an OD600 of 0.5–0.6, when 0.2% of arabinose was added and incubated for an additional of 2 hours at 37 °C. *ii*) followed by a second culture stage, in which the bacteria were transferred to LB medium supplemented with 0.5 mM IPTG and incubated for 18 hours at 20 °C.

To purify the Spy0128 protein co-expressed with the peptides, we followed the previously described protocol. Briefly, *E. coli* (BL21 pLysS) recently cultured were lysed by sonication in phosphate buffer (50 mM Na_2_HPO_4_/NaH_2_PO_4_, pH 7.0, 300 mM NaCl) supplemented with with 0.1 mM phenylmethylsulfonyl fluoride and 1 μM leupeptin. After centrifugation at 15,000g for 45 minutes, protein lysate was applied to Ni-NTA chromatography, washed three column volumes, and eluted with a nonlinear gradient of 250 mM imidazole diluted in phosphate buffer. We analyzed the protein fractions by SDS-PAGE and tracked them using Alexa488 HaloTag ligand, Alexa488 SNAP-Tag ligand, or Coomassie blue. The protein gels were imaged in a documentation gel station (Major Science, UVCI-1100) using epi illumination and proper filters. Selected Spy0128 polyprotein fractions were pooled and concentrated through Amicon filter units until reaching protein concentrations close to 10 μM, enough for the following step, biotinylation of the AviTag. Because phosphate buffer could interfere with the reaction with BirA, after concentrating the protein to 70-80 uL, the protein solution was diluted in 50 mM bicine buffer, pH 8,3, 10 mM ATP, 10 mM Mg(CH_3_COO)_2,_ and 50 μM biotin. The biotin reaction was initiated by adding 3 μL of 1mg/mL BirA and incubated either 3 h at 30 °C or 12 hours at 4 °C. After completing this last step, the protein was stored at −80°C after being supplemented with 20% glycerol. The protein is stable for weeks to months under these conditions.

### Surface Chemistry and Single Molecule Assays

For conducting the single-molecule force spectroscopy assay, we prepared custom-made fluid cells, similar to the procedure described previously. First, top and bottom cover glass were washed with 1% Hellmanex for 30 min in an ultrasonic bath at 50 °C and washed ten times with distilled water. The top glass, which is a 22 x 22 mm coverslip (n° 1, Ted Pella), was derivatized with repeal silane, making it completely hydrophobic. After incubating in the reagent for 1 hour at room temperature, we washed the top glass in 100% ethanol, dried it with compressed air, followed by a curing step at >60 °C for 1–2 h. After completing the modification, we transferred the top glass into a dissector for long-term storage. The bottom glass (24 x 60 mm, n° 1, Ted Pella) was washed with detergent and water, then transferred to a 1% solution of (3-aminopropyl) trimethoxysilane in ethanol and incubated for at least 20 minutes at room temperature. Then, after completing the silanization of the glass, we applied three washes with 100% ethanol, dried with compressed air, and cured for 1-2 hours at 100 °C. Similar to the top glass, we transferred the bottom glass to a dissector for long-term storage.

We assembled the fluid cells by sandwiching the top and bottom glass, separated by a thin layer of Parafilm, yielding a chamber of 300-500 µm high, two wells, and enough room to accommodate 30 μL of solution. To seal the fluid chamber, we melted the Parafilm in a heating block at 90°C and pressed the top glass onto the bottom with precision flat tweezers. Although the fluid chambers can be stored in a desiccator for weeks, we normally continue with the surface functionalization. Briefly, we added 1% glutaraldehyde diluted in PBS and incubated for 1 h, followed by adding an aliquot of 3.0 μm amine-functionalized polystyrene beads (Spherotech). These beads are later used in the experiments as fiduciary beads, allowing us to discount mechanical drift in the assay. After 20 minutes of bead incubation, we washed with PBS, eliminating unbound beads, and prepared the fluid chamber for the following step. An aliquot of 100 μL of amine O4 Halo-ligand (Promega) was added and incubated for at least 4 hours at room temperature, or overnight at 4 °C. After completing this last functionalization step, we washed the fluid chamber with PBS, considering at least a buffer change of 10× the fluid cell volume (300 μL). For blocking free amines available on the surface, we added a solution of 20 mM Tris-HCl pH 7.4, 150 mM NaCl, 2 mM MgCl_2_, supplemented with 1% BSA. The blocking reaction was left to react for at least 4 hours; to ensure that the buffer change occurred, we added a total of 200-300 μL of blocking solution. From here, the fluid chambers can be stored for one or two weeks at 4 °C.

### Protein Sequencing

A total of six gel-band samples were subjected to in-gel tryptic digestion followed by nano-liquid chromatography–tandem mass spectrometry (nLC–MS/MS). Excised bands were cut into approximately 1 mm³ pieces, washed sequentially with 25 mM ammonium bicarbonate and acetonitrile/water, and dried under vacuum. Proteins were reduced with 10 mM dithiothreitol in ammonium bicarbonate containing 10% acetonitrile at 56 °C, followed by alkylation with 55 mM iodoacetamide in aqueous ammonium bicarbonate for 45 min in the dark. Following additional washing and vacuum drying, proteins were digested with trypsin at 12.5 ng/µL in 25 mM ammonium bicarbonate for 6–8 h at 37°C. Peptides were recovered by sequential extraction with water and a solution containing 50% acetonitrile, 45% water, and 5% formic acid. Extracts were pooled, concentrated to approximately 10 µL using a vacuum centrifuge, and quantified using a NanoDrop instrument (Thermo Fisher Scientific). An aliquot containing 40 ng of peptides was loaded onto an Evotip (Evosep Biosystems).

Peptides were separated using an Evosep One nanoUHPLC system coupled to a timsTOF Pro 2 mass spectrometer (Bruker Daltonics). Chromatographic separation was performed on a PepSep C18 column (150 × 0.075 mm, 1.5 µm particle size) using the Whisper Zoom 40 samples-per-day method and a 36-min gradient from 2% to 35% solvent B, consisting of acetonitrile containing 0.1% formic acid. Data were acquired in data-dependent acquisition mode using timsControl 2.0 software and a PASEF method comprising 10 cycles. The acquisition range was m/z 100–1,700, with a capillary voltage of 1,500 V, a source temperature of 180°C, a time-of-flight repetition frequency of 10 kHz, and a resolving power of 50,000 FWHM.

Mass spectrometry data were analyzed using PEAKS Studio X+ (Bioinformatics Solutions), including the PEAKS IMS protein-identification workflow. Database searches were conducted against a combined UniProt database containing entries from Homo sapiens (9,606 proteins), *Streptococcus pyogenes* serotype M1 (1,690 proteins), and *Escherichia coli K-12* (4,402 proteins), the latter as a background for protein expression. Monoisotopic precursor and fragment mass tolerances were set to 50 ppm and 0.05 Da, respectively. Searches assumed specific tryptic digestion and allowed up to two missed cleavages per peptide. Cysteine carbamidomethylation was specified as a fixed modification. Additional modifications considered were methionine oxidation, lysine acetylation, asparagine and glutamine deamidation, and carbamylation of lysine residues and peptide N-termini. False discovery rates were estimated using a decoy database. Identifications were filtered using a PEAKS significance score threshold of ≥20, and protein identification required at least one unique peptide.

### Magnetic tweezers Assays and Data Interpretation

Before conducting the single-molecule experiments, we replaced the storage blocking buffer with a fresh buffer of 10 mM HEPES pH 7.8, 10 mM NaCl, supplemented with 1% BSA. We then added an aliquot of 30-50 μL containing 1-10 nM Spy0128 protein and allowed it to react for at least 15 minutes. After the reaction time, we washed the fluid chamber three times with the same Hepes buffer. Paramagnetic Dynabeads M270 (Invitrogen), previously prepared and blocked for at least 1 hour in the same Hepes buffer, were added to the fluid chamber and incubated for 5 to 15 minutes. An aliquot of 30-50 μL is enough to make the tethered beads available in the fluid chamber.

We acquired recordings using a laboratory-built magnetic tweezers microscope based on previous instrumentation descriptions. Our microscope is based on a 3D-printed inverted microscope, equipped with a piezoelectric and a linear focus actuator for moving the objective and a pair of permanent magnets. The custom-made software, also based on previous publications, allows us to track both beads: the reference polystyrene bead and the paramagnetic bead. We programmed magnet motion to vary the applied force while recording the position of the paramagnetic bead.

Because Spy0128 is inextensible in the absence of peptides, we selected only traces showing the presence of the I27 titin domains—25 nm in extension. By recording I27 unfolding, we are sure that the force traveled along the tethered protein molecule. Moreover, because we initially didn’t know the force at which the Spy0128-decorated isopeptide blocker unfolds, we used a force ramp from 4-100 pN, allowing us to record Spy0128 and the I27 titin domains in the same force pulse. Extension records and bead-position histograms were analyzed in Igor 8 (WaveMetrics).

## Results and Discussion

### Computational Isopeptide-blocker Design

During the inspection of the crystal structure of Spy0128, we concluded that a 10-residue peptide should be more than enough to establish a good match and form an intermolecular isopeptide bond. Thus, for building our peptide library, we consider a peptide of 10 residues long, where each position can be assayed by the 20 amino acids, excluding proline, which is known to present a low frequency in β-strands. Moreover, we held the ninth position as Asn, a residue needed for isopeptide bond formation. Thus, using our workflow, RPG reported a peptide library of more than 64 million peptides, from which we initially assayed a small subset of peptides, translated them to three-dimensional coordinates, and predicted binding energy through molecular docking. In the first iteration—termed *round A*—we converted more than 1,000 peptides to PDB files, docked them, and learned residue frequency across the 9 substituted positions. In the second iteration—*round B*—, we limited the 20 amino acid possibilities to high-frequency residues and eliminated amino acids with low or no presence in the previously docked peptides, yielding a library of 780k peptides. Finally, in a third iteration—*round C*—, using the biased amino acid distribution learned from the previous two iterations, we recovered a library of 350k peptides. After each iteration, we identified peptides with higher binding energy than the N14 peptide, previously reported as an effective isopeptide-blocker peptide. Peptide 96, from the third iteration, reached a docking score of −9.0 kcal/mol, more than 1 kcal/mol more stable than the N14 peptide (see **table/Figure**). The peptide, now termed C96 and with the sequence *QWGNMFNKWK*, showed in the docking assay to display an optimal orientation, locating the side chain of Asn4 facing Lys168 (**Figure** 2).

**Figure 2.**
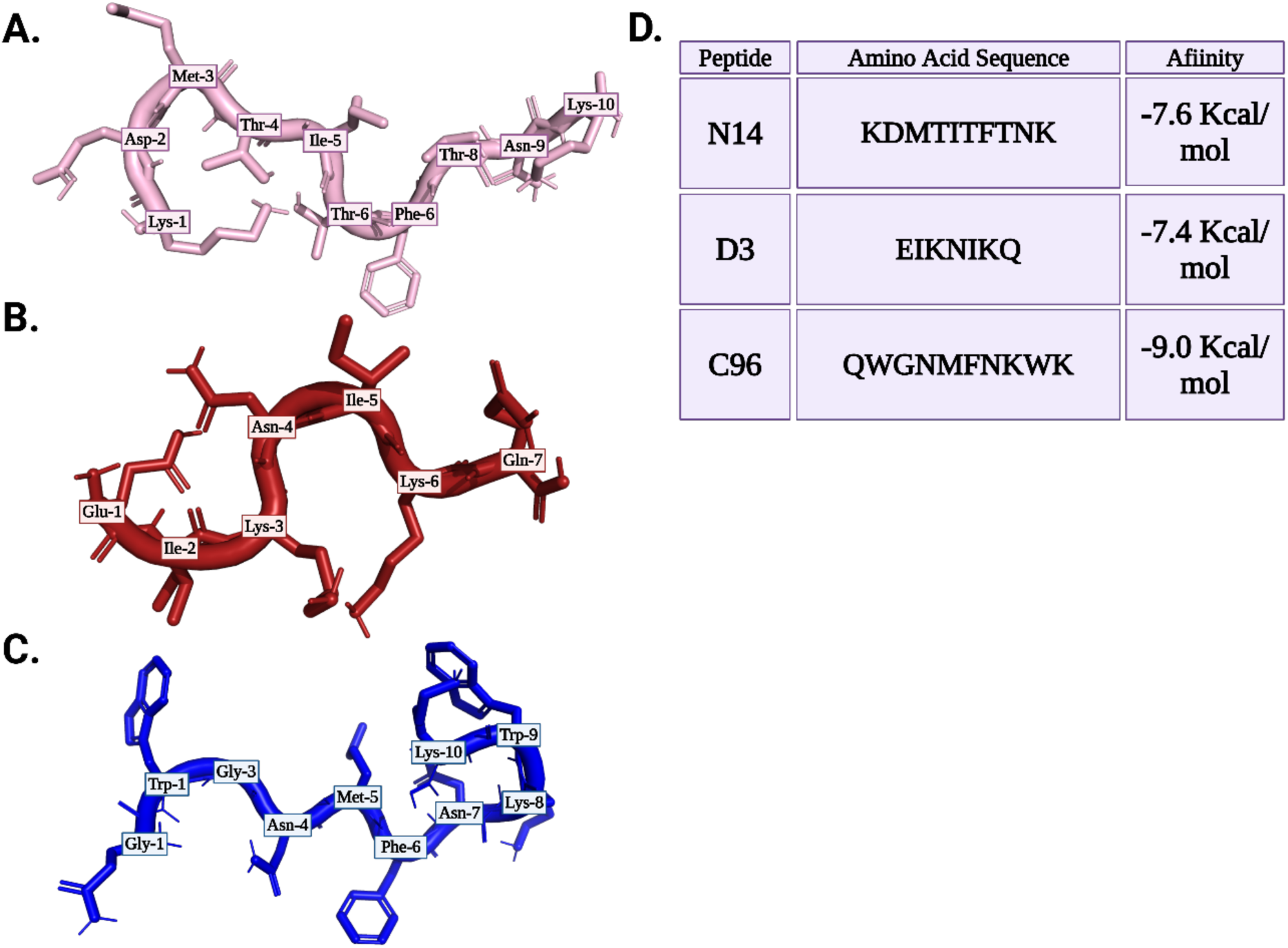
Peptide models and docking scores obtained with RPG A–C. Docking representations associated with the N14 reference, D3, and C96, respectively. A shortened β11-derived representation was used for computational comparison with N14 [4]; the experimental N14 blocker is the peptide described in the earlier study [1]. D. Docking scores were −7.6 kcal/mol for the N14 reference model, −7.4 kcal/mol for D3, and −9.0 kcal/mol for C96. The more favorable score of C96 motivated its selection for biochemical and mechanical evaluation. These scores rank the calculated arrangements and are not experimental binding affinities.

In addition to the N14 and C96 peptides, we computationally assayed several peptides: randomized sequences, Ala- or Gly-based sequences, and a peptide originally designed to target a different pili protein—SpaD from *Corynebacterium diphtheriae.* All these peptides show less binding energy compared to N14 and C96 peptides. Thus, to prove in the *in vitro* assay the role of the mimicking design –N14 peptide–and computational prediction –C96 peptide, we decided to use the peptide D3, designed for *C. diphtheriae*.

### Protein Co-expression and Isopeptide interfering

To confirm that both vectors contain either the Spy028 polyprotein or the peptide-tagged protein, we tested them by individual induction. As expected, the Spy0128 polyprotein migrates near 120 kDa, while the SNAP-tagged peptides are below 20 kDa, consistent with the theoretical molecular weight (**Figure**). Thus, with the individual expression confirmed, we continued with the sequential expression—induction with arabinose for triggering the production of the SNAP-tagged peptide, followed by IPTG for Spy0128 polypeptide—. Using fluorescent ligands, we confirmed in cell lysates the presence of the engineered Spy0128 polyprotein, along with several other low-molecular-weight bands; however, we couldn’t confirm bands that account for the Spy0128 polyprotein bound to SNAP-tagged peptides. To better resolve Spy0128 decoration, we purified the co-expressed protein and analyzed it by SDS-PAGE, labeling each fraction. We detected a prominent band near 130 kDa in all three conditions, matching the unmodified target construct. We observed additional bands migrating in the upper region of the gels, with an apparent mass near 250 kDa, probably due to the formation of the Spy0128-SNAP-tagged peptide. These species were visible with HaloTag fluorescence and His-tag staining (**Figure 3C**). To get an overall picture of the three systems, we pooled the Ni-protein fractions and ran an SDS-PAGE with prelabeled Halo-ligand at a higher protein concentration. Surprisingly, InVision and HaloTag labeling also revealed bands near 100, 70, and 50 kDa. These bands are consistent with heterogeneity of the purified preparation and could arise from partial proteolysis. Although the construct architecture suggests possible fragments (**Figure 3D**), apparent molecular mass does not establish their sequence or cleavage site.

**Figure 3.**
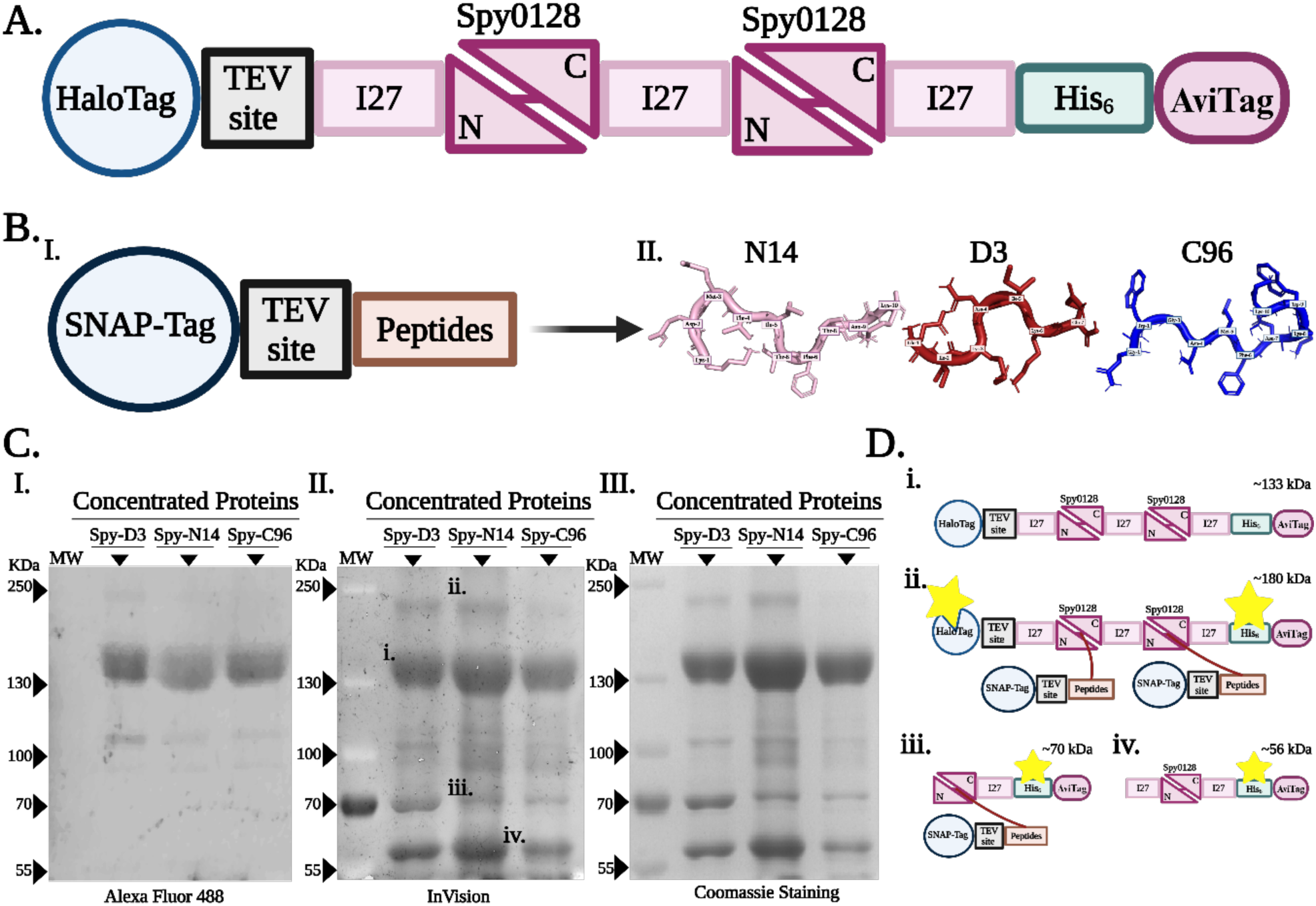
Constructs and electrophoretic analysis of peptide–Spy0128 coexpression products A. The target construct contains HaloTag, two Spy0128 repeats flanked by three I27 domains, a His-tag, and AviTag. B. Peptides N14, D3, and C96 were expressed as SNAP-tag fusions containing a TEV recognition site. C. Purified products were separated by 8% SDS-PAGE and detected using HaloTag-associated Alexa Fluor 488 fluorescence (I), InVision His-tag staining (II), and Coomassie staining (III). Lane order is D3, N14, and C96. A prominent target band near 130 kDa and additional higher-molecular-mass species are present across the conditions. Bands near 70 and 56 kDa indicate additional species in the preparation. Comparisons of intensity are qualitative. D. Schematics illustrate proposed intact, peptide-associated, and fragmented constructs. These assignments are hypotheses based on construct architecture, rather than molecular identities established by the gels. HaloTag and His-tag signals identify the target construct and do not independently establish incorporation of the SNAP-tagged peptide.

### Force Spectroscopy Reveals Spy0128 Unfolding

We used magnetic tweezers to test whether the peptide-associated species detected by electrophoresis also exhibited an altered mechanical extension. The Spy0128 polyprotein was tethered through HaloTag—bound directly to the glass surface— and biotinylated AviTag—recognized by the Streptavidin on the paramagnetic bead—(**Figure 4A**). We compared recordings obtained without peptide and after coexpression with the designed peptides D3, N14, or C96. In general, the four conditions show similar mechanical behavior; I27 titin domains unfold along the force ramp. Nevertheless, in some cases, the peptides yield extra events. Despite the low probability of these events, the extension may suggest Spy0128 unfolding. Comparing N14, D3, and C96 peptide coexpression, the abundance of trajectories with extra events does not differ much, probably because, despite the intervention’s success, it remains a low-probability event.

**Figure 4.**
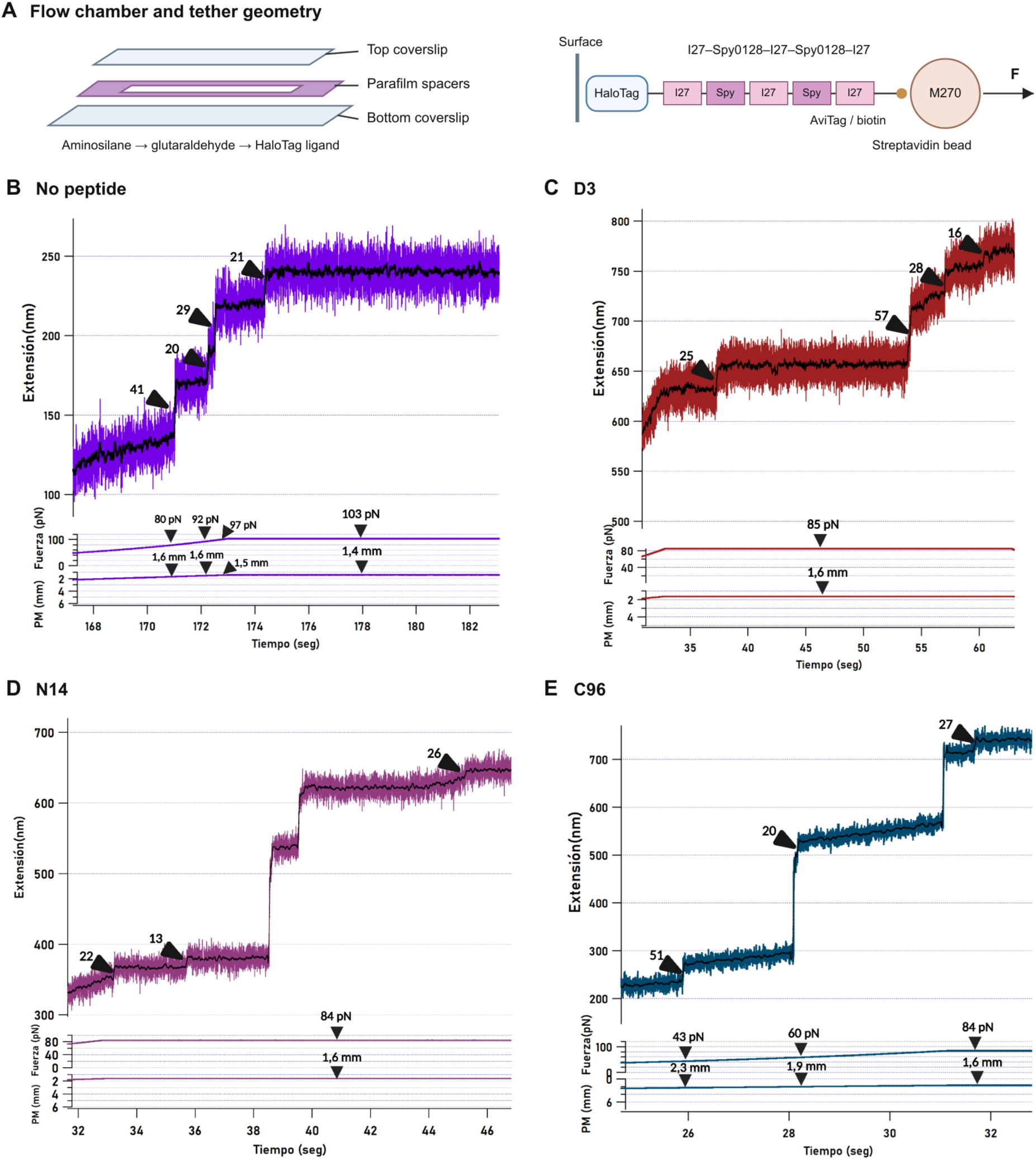
Magnetic tweezers recordings of Spy0128 with and without peptides Flow chamber and HaloTag–AviTag tether geometry. B–E. Original extension recordings without peptide (B), with D3 (C), with N14 (D), and with C96 (E), from thesis Figures 4.11, 4.13, 4.15, and 4.17 [4]. Step annotations are in nanometers. Force and magnet-position histories, time axes, and original extension coordinates are retained; axis ranges differ. The peptide-free record resolves three I27-like events. N14 and C96 each resolve two I27-like events together with additional extension. These changes could include extensible Spy0128, but the incomplete reporter fingerprints prevent unambiguous full-length assignment. Absolute extension levels are not contour-length increments or a measure of blocking efficiency. The original Spanish axis labels denote extension, force, magnet position (PM), and time.

### Mass Spectroscopy Confirmed Protein Hydrolysis

Given that the designed peptide interferes with isopeptide bond formation without increasing the amount of decorated Spy0128, similar to the control N14 peptide, we explored the idea that the designed peptide triggers protein hydrolysis. The protein degradation supports the idea that the peptides prevent Spy0128 from folding properly, allowing protein hydrolysis within the cell. To confirm this scenario, we sequenced the protein bands present in the N14 and C96. We examined the bands at 100, 75, and 50 kDa through mass spectrometry, confirming the presence of I27, HaloTag, and the designed C96 peptide. Therefore, it is likely that the peptide triggers protein hydrolysis after successfully preventing the formation of the intramolecular isopeptide bond and interfering with Spy0128 folding.

## Discussion and Conclusion

The original Spy0128 study established that a peptide could compete with native isopeptide bond formation during pilin folding and thereby weaken the resulting protein. Here, we extend the candidate-selection stage of that strategy. RPG provides a route from sequence generation to structural preparation and docking, while the expression system permits selected peptides to be produced before the target. C96 emerged with a more favorable docking score than the N14 reference model. Electrophoresis and magnetic tweezers then provided complementary observations: recovery of higher-molecular-mass target-containing species and recordings with extensions beyond the identifiable reporter pattern. Moreover, mass spectroscopy confirms that lower-molecular-mass bands are hydrolyzed Spy0128 protein.

This distinction is central to interpreting the computational results. A docking score evaluates a modeled peptide–protein arrangement. It does not determine whether the peptide can encounter a nascent pilin, compete with its native β-strand, or support an intermolecular isopeptide reaction before the intramolecular bond forms. The earlier study showed that this timing is essential [1]. A favorable score against a prepared target structure therefore provides a selection criterion, rather than evidence that the folding-dependent intervention has occurred.

The biochemical readout also requires a different interpretation from that used in the original study. With HaloTag on the peptide, fluorescent labeling of a higher-molecular-mass Spy0128 species supported recruitment of the tagged blocker [1]. With HaloTag on the target, fluorescence remains present whether or not a peptide is incorporated. InVision detects a second tag on that same target construct and therefore corroborates target identity rather than peptide identity. The additional species could represent peptide-associated products, but oligomerization or other changes in the purified construct remain alternative explanations. Direct detection of the SNAP-tagged peptide in an isolated species would provide a more discriminating test.

The D3 result further limits a specificity claim. D3 was designed for domain 3 of SpaD from Corynebacterium diphtheriae, yet its coexpression condition produced a prominent upper band. This observation could reflect cross-reactivity or a process unrelated to the intended Lys179-directed reaction. It does not establish either explanation. Target-only, tag-only, and reversed-induction controls, together with direct characterization of the recovered species, would help distinguish them. Comparison of docking scores with quantified modification yields would then test whether computational ranking predicts the intended biochemical outcome.

The magnetic tweezers experiments address the mechanical consequence of peptide coexpression directly. The peptide-free example displays three I27-like reporter events, whereas the N14 and C96 examples combine only two clearly identifiable I27-like steps with additional changes in extension. Such changes are compatible with extensible Spy0128, consistent with the mechanism established previously [1]. The difficulty of obtaining an unambiguous full-length fingerprint, however, prevents us from treating every long recording as an intact, peptide-modified polyprotein. In particular, a large absolute extension or an unassigned jump is not sufficient to identify unfolding of one Spy0128 domain. These records support the proposed interpretation qualitatively, while the completeness of the tether and the force dependence of each event remain important constraints.

The additional data also do not resolve the specificity question raised by D3. Its biochemical and mechanical behavior must be considered alongside N14 and C96 rather than excluded because of its intended role as a negative control. A larger set of full-length records, matched force protocols, and sequence-dependent controls would be needed to compare blocking efficiency. The current recordings do not establish a peptide-specific modification yield or an effect on bacterial adhesion or antimicrobial activity.

RPG therefore connects candidate selection with expression, biochemical characterization, and an initial mechanical test. Among the peptide-associated tethers that could be followed, some extensions were not explained solely by the recognizable I27 reporters and could include extended Spy0128. The main experimental limitation is reliably identifying complete constructs. Resolving that limitation and establishing the chemical identity of the peptide-associated species will determine how effectively computationally selected peptides interfere with native isopeptide bond formation.

**Table 1.**
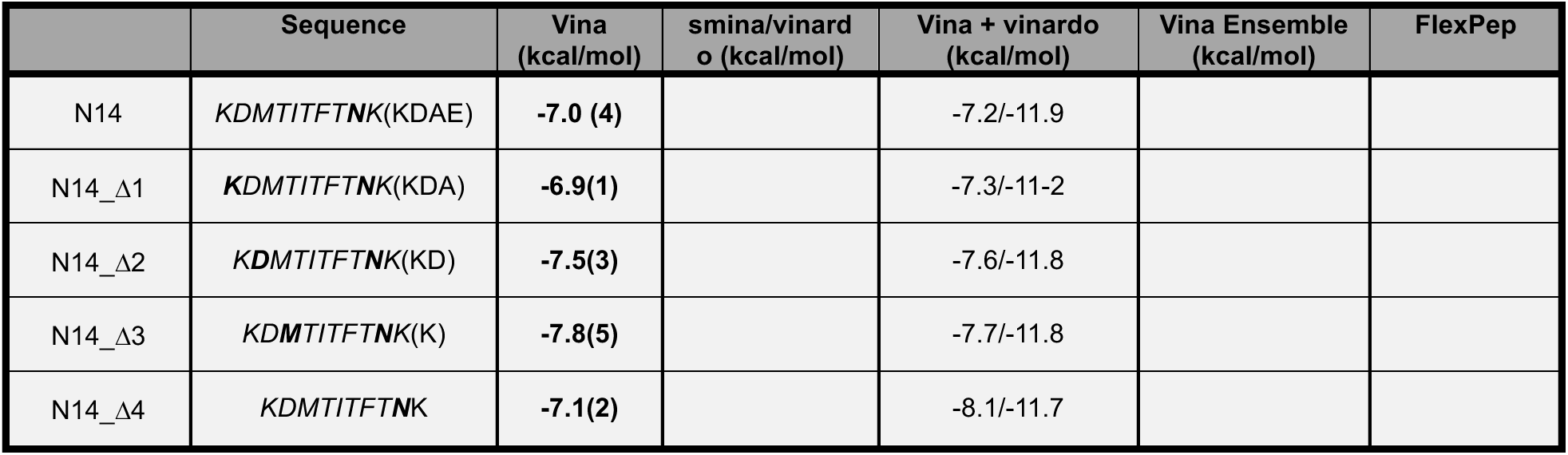
Binding energy of 10-amino-acid length.

|  | **Sequence** | **Vina (kcal/mol)** | **smina/vinardo (kcal/mol)** | **Vina + vinardo (kcal/mol)** | **Vina Ensemble (kcal/mol)** | **FlexPep** |
|---|---|---|---|---|---|---|
| N14 | *KDMTITFT**N**K*(KDAE) | **-7.0 (4)** |  | -7.2/-11.9 |  |  |
| N14_∆1 | ***K**DMTITFT**N**K*(KDA) | **-6.9(1)** |  | -7.3/-11-2 |  |  |
| N14_∆2 | *K**D**MTITFT**N**K*(KD) | **-7.5(3)** |  | -7.6/-11.8 |  |  |
| N14_∆3 | *KD**M**TITFT**N**K*(K) | **-7.8(5)** |  | -7.7/-11.8 |  |  |
| N14_∆4 | *KDMTITFT* ***N***K | **-7.1(2)** |  | -8.1/-11.7 |  |  |

**Table 2.** Determination of critical position along the N14 peptide.

|  | **Sequence** | **Vina (kcal/mol)** | **smina/vinardo (kcal/mol)** | **Vina + vinardo (kcal/mol)** | **Vina Ensemble (kcal/mol)** | **FlexPep** |
|---|---|---|---|---|---|---|
| N14 | *KDMTITFT**N**K*(KDAE) | -7.7 | -10.3 | -7.6 / -10.9 | -7.7/-7.6 |  |
| N14_Ala1 | ***A**DMTITFT**N**K*(KDAE) | -7.8 | **-8.3** | **-8.0 / -11.0** | **-8.1/-8.3** |  |
| N14_Ala2 | *K**A**MTITFT**N**K*(KDAE) | -7.5 | -10 | -7.9 / -11.2 | **-8.0/-8.1** |  |
| N14_Ala3 | *KD**A**TITFT**N**K*(KDAE) | **-8.1** | **-8.1** | -7.5 / -10.8 | **-8.0/-8.0** |  |
| N14_Ala4 | *KDM**A**ITFT**N**K*(KDAE) | -7.9 | -9.7 | -7.9 / -11.8 | -7.9/-7.9 |  |
| N14_Ala5 | *KDMT**A**TFT**N**K*(KDAE) | -7.8 | **-6.8** | **-8.0 / - 11.2** | -7.4/-7.4 |  |
| N14_Ala6 | *KDMTI**A**FT**N**K*(KDAE) | **-8.6** | -7.8 | -7.9 / -11.2 | -7.5/-7.7 |  |
| N14_Ala7 | *KDMTIT**A**T**N**K*(KDAE) | -7-4 | **-5.2** | -7.7 / -11.5 | -7.2/-7.8 |  |
| N14_Ala8 | *KDMTITF**AN**K*(KDAE) | **-6.8** | -10.1 | -7.0 / -11.3 | -7.7/-7.8 |  |
| N14_Ala9 | *KDMTITFT**A**K*(KDAE) | -7.2 | **-7.0** | **-8.0 / - 11.2** | -7.8/-7.8 |  |
| N14_Ala10 | *KDMTITFT**NA***(KDAE) | -7.6 | -9.8 | -7.6 / -10.8 | **-8.1/-8.7** |  |

**Table 3.** Binding energy of the engineered peptides.

|  | **Sequence** | **Vina<br>(kcal/mol)** | **smina/vinardo<br>(kcal/mol)** | **Vina + vinardo<br>(kcal/mol)** | **Vina Ensemble<br>(kcal/mol)** | **FlexPep** |
|---|---|---|---|---|---|---|
| N14_Spy | *KDMTITFT**N**K*(KDAE) | -7.3 | -12.2 | -7.4/-10.9** | -7.7/-8.7*** |  |
| D3_Cdiph | *EIK**N**IKQ* | -7.9 | -8.51 | -7.5/-10.4** | -7.6/-7.6*** |  |
| Ala10 | *AAAAAA**A**AAA* | -8.1 | -5.0 | -8.6/-9.9** | -8.6/ -8.6*** |  |
| N14_Ala | *AAAAAA**N**AAA* | -7.5 | -5.8! | -8.5/-9.4** | -8.3/-7.6*** |  |
| N14_Gly | *GGGGGG**N**GGG* | -8.2 | -7.3! | -7.0/-7.4** | -7.6/-8.3*** |  |
| C96 | *QWGNMF**N**KWK* | -9.1 | -6.2 | -8.3/-11.6** | -8.1/-8.4*** |  |

\*\* vinardo score; \*\*\* emsable score

## Acknoledgments

This work was supported by ANID FONDECYT grant 1221064. We also acknowledge the funds from Universidad Mayor FDP2021-20 to support VC research. We thank all members of the Rivas Pardo laboratory for their helpful discussions.

## Author contributions

J.A.R.P. conceived the project. M.M., T.H., and J.A.R.P. designed the experiments, while T.H. and M.M. conducted the RPG workflow and peptide design. M.M. and V.C. conducted the *in vitro* experiments, whereas M.M. examined the protein using single-molecule force spectroscopy. J.A.R.P. and M.M. drafted the paper with input from all authors.

## Additional information

Supplementary information is available for this paper at www.journal.com

